# Serum from pregnant donors induces human beta cell proliferation and insulin secretion

**DOI:** 10.1101/2023.04.17.537214

**Authors:** Kendra R. Sylvester-Armstrong, Callie F. Reeder, Andrece Powell, Matthew W. Becker, D. Walker Hagan, Jing Chen, Clayton E. Mathews, Clive H. Wasserfall, Mark A. Atkinson, Robert Egerman, Edward A. Phelps

## Abstract

Pancreatic beta cells are among the slowest replicating cells in the human body. Human beta cells usually do not increase in number with exceptions being during the neonatal period, in cases of obesity, and during pregnancy. This project explored maternal serum for stimulatory potential on human beta cell proliferation and insulin output. Gravid, full-term women who were scheduled to undergo cesarean delivery were recruited for this study. A human beta cell line was cultured in media supplemented with serum from pregnant and non-pregnant donors and assessed for differences in proliferation and insulin secretion. A subset of pregnant donor sera induced significant increases in beta cell proliferation and insulin secretion. Pooled serum from pregnant donors also increased proliferation in primary human beta cells but not primary human hepatocytes indicating a cell-type specific effect. This study suggests stimulatory factors in human serum during pregnancy could provide a novel approach for human beta cell expansion.

## INTRODUCTION

Glucose metabolism changes during pregnancy to support the growing fetus’ needs for glucose through passive transport dependent on a concentration gradient between fetal and maternal circulations. In the later stages of pregnancy, the placenta secretes hormones to increase maternal insulin resistance and hepatic glucose production, raising the maternal glucose, supporting the gradient [1]. To prevent excessive nutrient delivery to the fetus and prevent maternal hyperglycemia, pregnancy is also associated with an increase in beta cell mass [1, 2]. This increase in beta cell mass during human pregnancy has been investigated by two autopsy series [3, 4]. However, the dominant mechanism (replication, neogenesis or both) for increased beta cell mass during pregnancy in humans remains uncertain [1, 4, 5].

In rodents, pregnancy-associated beta cell expansion depends on secreted placental lactogens that signal through the prolactin receptor [2]. At the end of pregnancy, the beta cell population contracts back to its pre-pregnancy size. However, loss of prolactin associated signaling pathways in the mouse islet does not completely block the proliferative response to pregnancy. Studies of lactogens on human beta cell proliferation have generated inconclusive results [6, 7]. Therefore, other signals almost certainly contribute to beta cell proliferation in pregnancy [5]. Serotonin has been identified as one such signal that regulates beta cell mass and that is induced during pregnancy [8-10].

## MATERIALS AND METHODS

### Study Subjects and Sample Collection

Gravid, full-term women who were scheduled to undergo cesarean delivery were recruited at the Department of Obstetrics and Gynecology at the University of Florida from September 2020 to October 2020 for serum sample collection. Women were dated by first trimester ultrasound and prenatal ultrasound examination confirmed that there were no anatomical abnormalities of the fetuses. All participants underwent glucose tolerance testing from 24-28 weeks and were confirmed to have normal glucose tolerance, gestational or pre-gestational diabetes.

### Statistical analysis

All data are presented as the mean ± SEM. The statistical significance was calculated by Student’s t test (two tailed) or analysis of variance (ANOVA) followed by post-hoc pairwise comparisons. Statistical significance was defined as at least p<0.05. The analysis was performed in GraphPad Prism version 9.0.

### Study Subjects and Sample Collection

Study data were collected and managed using REDCap electronic data capture tools hosted at the University of Florida. After written, informed consent for participation was obtained, maternal peripheral blood samples were collected just prior to delivery. Centrifugation was performed to obtain serum and plasma samples (2000 x g at 4°C for 15 min). Serum and plasma samples were stored at -80°C until analyzed. Baseline blood glucose and insulin concentrations were measured on all maternal serum samples using an Accu-Check Aviva handheld glucose meter and human insulin Chemiluminescence ELISA kit (ALPCO).

### EndoC-βH1 beta cell culture

The EndoC-βH1 human beta cell line was cultured consistent with conditions described by Tsonkova et al [11]. The EndoC-βH1 was cultured in DMEM low glucose (1g/L), 2% Albumin from bovine serum fraction V, 50 μM 2-mercaptoethanol, 10 mM nicotinamide, 5.5 μg/ml transferrin, 6.7 ng/ml sodium selenite and Penicillin (100 units/ml) / Streptomycin (100 μg/ml) and maintained in sub confluent densities. EndoC-βH1 cells were seeded at 2 × 10^5^ cells per well in a cell culture treaded 96-well plate and allowed to adhere for 24 hours prior to starting of the experiments. The next day, seeding media was exchanged for fresh media containing 10% human serum from pregnant or non-pregnant donors together with EdU (Click-iT™ EdU Proliferation Assay for Microplates, ThermoFisher Scientific) and cultured for 1 week. At takedown, EdU was detected following the kit manufacturer’s protocol and fluorescence was measured with a microplate reader (Molecular Dives Spectramax M5) at 568/585 nm. Secreted insulin in cell culture supernatant was measured by human Insulin Chemiluminescence ELISA kit (ALPCO).

### Primary Human Beta Cells

A detailed description and validation of monolayer cultures of primary human and rat islets cells is available [12]. Primary human pancreatic islets from deceased nondiabetic donors were obtained from the Integrated Islet Distribution Program (IIDP) at City of Hope. Human islets were cultured at 24°C and 5% CO2 in 10 cm non-adherent cell culture dishes in CMRL medium with 2% L-glutamine, 10% fetal bovine serum (FBS), 10 mM HEPES and 1% penicillin/streptomycin. Islets were dissociated into a suspension of single islet cells by continuous gentle pipetting in 0.3 ml 0.05% trypsin-EDTA per 500 islets for 3 min at 37 °C. Islet single cells were seeded 35,000 cells per cm^2^ on coverslip bottom 96 well microplates (Ibidi).

Wells were precoated with purified human collagen IV (MilliporeSigma) at 50 μg/ml in HBSS with Ca^2+^/Mg^2+^ for 1 h at 37 °C. Islet cells required 3–4 d of culture to adhere and spread on surfaces before further experimentation. Islet monolayers were cultured in minimum essential medium (MEM) with GlutaMAX, 11 mM glucose, 5% FBS, 1 mM sodium pyruvate, 10 mM HEPES and 1× B-27 Supplement + 10% human serum. Once primary human islet monolayers were established, 10% pooled human serum from pregnant or non-pregnant donor groups was added together with EdU (Click-iT EdU Cull Proliferation kit for Imaging, Thermo Fisher Scientific) and cultured for two weeks. Samples from the primary human monolayer experiment were fixed and immunostained for insulin with guinea pig anti-insulin primary antibody (Dako A0564) and Alexa-Fluor 568 goat anti-guinea pig secondary antibody (Thermo Fisher Scientific), developed for EdU detection following the kit manufacturer’s protocol, and counterstained with DAPI. Samples were imaged on a Leica SP8 Confocal microscope with a 20x / 0.8 numerical aperture Plan-Apochromat air objective.

### Primary Human Hepatocytes

Cryopreserved primary human hepatocytes (Lonza #HUCPG) were cultured according to the supplier’s instructions. Hepatocytes were thawed at 37°C and placed in warmed thawing medium (Lonza # MCHT50P), resuspended in Hepatocyte Plating Medium (Lonza #MP100), plated in collagen-coated 96 well plates at 50,000 cells/well, and cultured at 37°C and 5% CO2. Culture wells were precoated with 140 μg/ml Collagen I solution (Roche #11179179001) for 2 hrs at 37°C. The initial plating medium was exchanged for fresh plating medium after 1h. Plating media was then exchanged again for Hepatocyte Culture Medium (Lonza #CC-4182) after 24 h, and then daily for the next 3 days. Once primary human hepatocyte monolayers were established, 10% pooled human serum from pregnant or non-pregnant donors was added together with EdU (Click-iT™ EdU Proliferation Assay for Microplates, ThermoFisher Scientific) and cultured for one week. At takedown, EdU was detected following the manufacturer’s protocol and fluorescence was measured with a microplate reader (Molecular Dives Spectramax M5) at 568/585 nm.

## RESULTS

### Participant characteristics

A total of 20 pregnant women were recruited for this study. Clinical characteristics of the donor group are detailed in **Table 1 and Supplementary Table 1**. Serum samples from 15 of the donors were selected for further experiments and 5 of the donor samples were excluded due to hemolysis. All women in this study underwent scheduled cesarean delivery with a full-term, non-anomalous singleton gestation without intrauterine infection or other obstetric complications. 17 participants had normal glucose tolerance, two had gestational diabetes and one had type 2 diabetes. Mean maternal age was 31 ± 5.52 years and BMI averaged 35.90 ± 8.09 kg/m^2^. The mean gestational age at delivery was 39 ± 0.79 weeks. Banked sera from an additional 6 pregnant donors was selected to repeat the experiment.

**Table 1.** Clinical Characteristics of Patients and Neonates.

| Characteristic | (n=20) |
| --- | --- |
| Maternal Variables |  |
| Age (years) | 30.2 ± 4.7 (19-37) |
| Race |  |
| White | 15 (65%) |
| Black | 4 (20%) |
| Hispanic | 2 (10%) |
| Other | 1 (5%) |
| BMI (kg/m<sup>2</sup>) | 35.90 ± 8.09 (26.9-53.9) |
| First trimester A1C<sup>a</sup> | 5.70 ± 0.37 (5.3-6.18) |
| Gestational age at delivery (weeks) | 39 ± 0.79 (37 – 40) |
| Glucose Tolerance |  |
| Normal glucose tolerant | 17 |
| Gestational diabetes mellitus | 2 |
| Type 1 diabetes mellitus | 0 |
| Type 2 diabetes mellitus | 1 |
| Newborn Variables |  |
| Placental weight (g) | 650 ± 157.56 (497-905) |
| Neonatal weight (g) | 3450 ± 390.38 (2992–4250) |
| Fetal sex (male/female) | 13/7 |

<sup>a</sup>Not all donors had first trimester hemoglobin A1Cs drawn. Thus, mean is based off n=5 donors

### Effect of maternal serum on EndoC-βH1 beta cells

We measured a donor-dependent effect of supplementing the cell culture medium with 10% maternal serum on EndoC-βH1 beta cell proliferation by EdU incorporation, and on insulin secretion by ELISA (**Figure 1**). There were no significant differences in the degree of proliferation for EndoC-βH1 beta cells cultured with 10% maternal serum, 10% non-pregnant serum, or serum-free control medium when samples were analyzed by treatment group (p=0.78) (**Figure 1A**). There was, however, a distinct group of maternal “responders” (n=4; donors #5-8), which induced a higher degree of proliferation relative to control medium. When analyzed as responders and non-responders, the proliferation induced by these four donors was statistically significant (p=0.003). (**Figure 1B and Supplementary Figure 1**). Responders were selected as having a mean more than 1 standard deviation higher than the mean of the standard cell culture medium control group.

**Figure 1.**
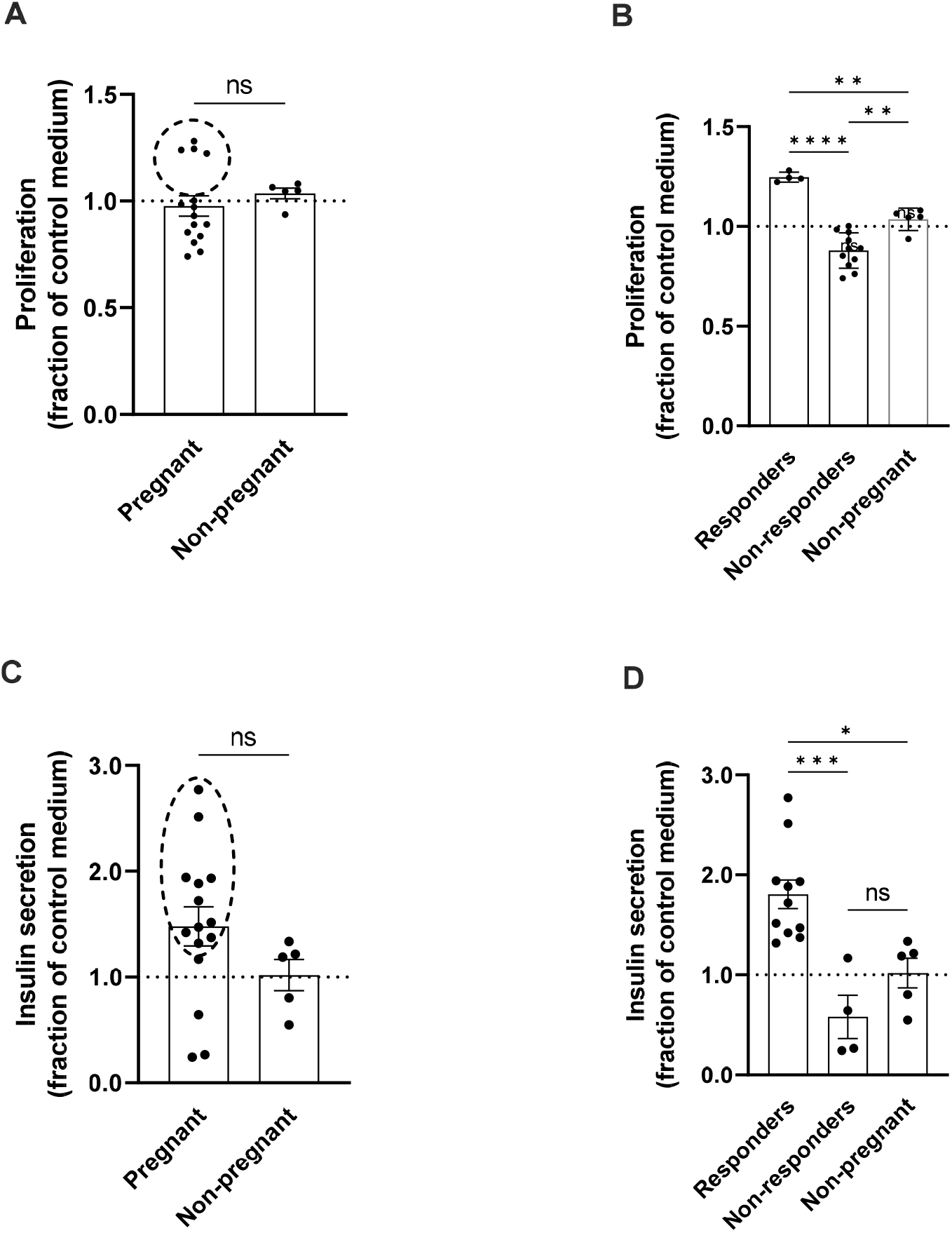
Maternal serum effects on EndoC beta cell proliferation and insulin secretion. EndoC beta cells were cultured in 96-well plates supplemented with 10% pregnant or non-pregnant donor serum or control medium (serum-free defined growth medium) for seven days. **A-B**. Total EndoC beta cell proliferation, normalized relative to control medium, measured by EdU incorporation. **(A)** Pregnant (n=15 donors) v. non-pregnant (n=5 donors) or **(B)** Pregnant donors subdivided into responders (n=4 donors) and non-responders (n=11 donors) v. non-pregnant (n=5 donors). **C-D**. Accumulated EndoC beta cell insulin secretion to culture medium, normalized relative to control medium, measured by ELISA kit. **(C)** Pregnant (n=15 donors) v. non-pregnant (n=5 donors) or **(D)** Pregnant donors subdivided into responders (n=11 donors) and non-responders (n=4 donors) v. non-pregnant (n=5 donors). Groups were assessed for differences by two-tailed t test (A,C) or one-way ANOVA with Tukey’s post hoc pairwise comparisons (B,D). Each dot represents the mean proliferation response (A,B) or concentration of secreted insulin (C,D) of three technical replicates for an individual donor as a fraction of the mean response to control serum-free medium. Values are mean ± SEM. *p<0.05, **p<0.01, ***p<0.001,****p<0.0001.

A different group of maternal “responders” (n=12: donors 1-12) within the maternal serum group, induced significantly increased insulin secretion (p=0.02) to the media over 24 hours of culture in 5.5 mM glucose, relative to control medium (**Figure 1D and Supplementary Figure 1)**. There was no correlation of proliferation with insulin secretion. In other words, the donors that induced higher levels of proliferation were not the same donors that induced higher levels of insulin secretion.

We repeated the proliferation experiment with a different group of six pregnant donors (**Figure 2**) collected during the third trimester and 24-48 hours post-delivery. In the second experiment, three out of six pregnant donors significantly increased EndoC-βH1 beta cell proliferation while FBS or serum from male donors did not. Blood collected during the third trimester and post-partum both imparted similar degrees of beta cell proliferation and results were consistent between donors so these datapoints were pooled together (**Figure 2**).

**Figure 2.**
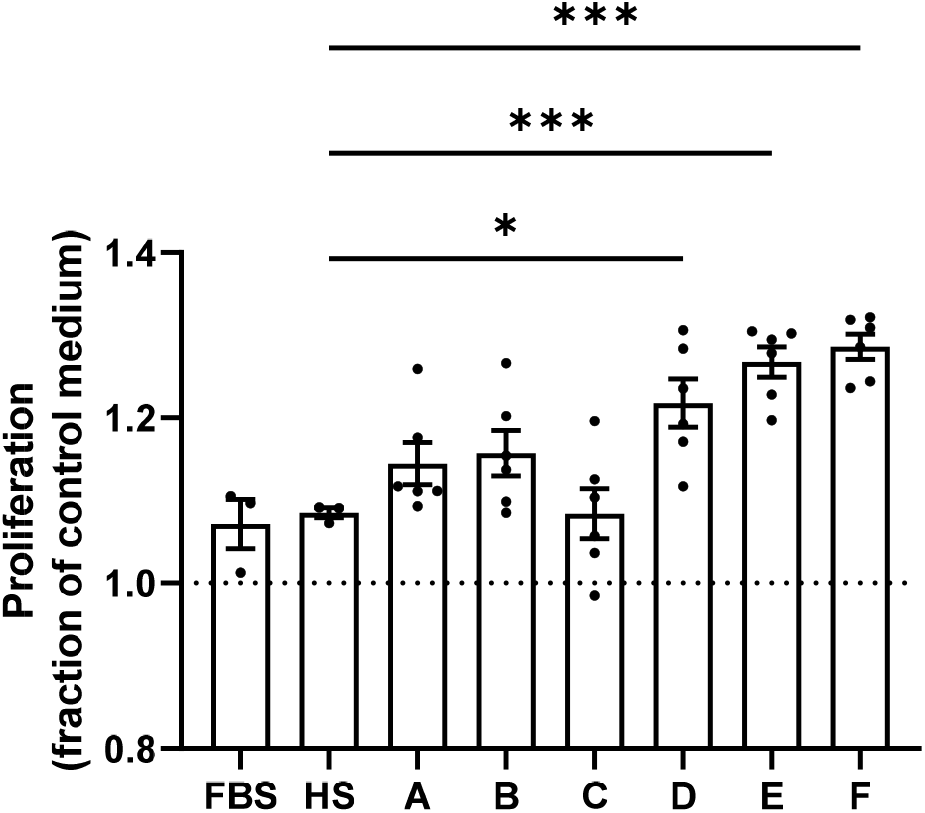
Second set of pregnant human serum donors showing donor-specific increase in EndoC beta cell proliferation compared to serum-free control medium. EndoC beta cells were cultured in 96-well plates supplemented with 10% pregnant human serum (Donors A-F) or pooled male, non-pregnant human serum (HS) or 10% fetal bovine serum (FBS) for seven days. Proliferation was measured by EdU incorporation. Groups were assessed for differences by one-way ANOVA with Tukey’s post hoc pairwise comparisons. Each dot represents the proliferation response in an individual well as a fraction of the mean response to control serum-free medium. Values are mean ± SEM. *p<0.05, ***p<0.001

In rodents, prolactin hormones have been identified as a mechanism of enhanced beta cell proliferation and insulin output during pregnancy [1, 2]. Differential prolactin levels could potentially explain the higher levels of proliferation or insulin secretion in our study observed for some, but not all, donors. We measured prolactin in our collection of maternal serum samples by ELISA and plotted it against beta cell proliferation and insulin secretion for each donor (**Supplementary Figure 2**). Unlike what has been reported for rodents, there was not a significant correlation between serum prolactin and our measurements of beta cell proliferation or insulin secretion, although the correlation with prolactin was stronger for proliferation than insulin secretion (r^2^ value of 0.20 versus 0.02). Perhaps a larger study cohort would reveal a modest correlation between prolactin levels and beta cell proliferation.

### Effect of maternal serum on primary human islets

Lastly, we tested the effect of maternal serum on monolayers of dispersed primary human islets to analyze proliferative effects of maternal serum. Monolayers of dispersed islet cells are a technique we and others have used to identify EdU^+^/insulin^+^ beta cells which are more difficult to accurately assess in three-dimensional whole islets [12-15]. After two weeks of incubation with pooled serum from pregnant and nonpregnant donors, human islet cell monolayers were fixed and immunostained for insulin and developed for EdU incorporation (**Figure 3A-D**). Because human islets are limited, we pooled serum from four pregnant donors and compared this to pooled serum from four non-pregnant donors, rather than attempt to identify individual donors as responders and non-responders. As is typical for primary human islets, cell proliferation rates were low across all samples, requiring confocal imaging to identify proliferating cells. However, a significantly higher percentage of Insulin^+^/EdU^+^ cells were observed for monolayers cultured with pregnant serum. This result was repeated in a second experiment using a different islet donor. The primary human beta cell proliferation rates of around 1% we observed with pregnant serum are similar with those observed for harmine [13]. To verify the effect was cell-type specific, primary human hepatocytes were similarly cultured with pooled pregnant and non-pregnant human serum, EdU incorporation was detected by microplate assay, and there was no differential effect on hepatocyte proliferation (**Figure 3E**).

**Figure 3.**
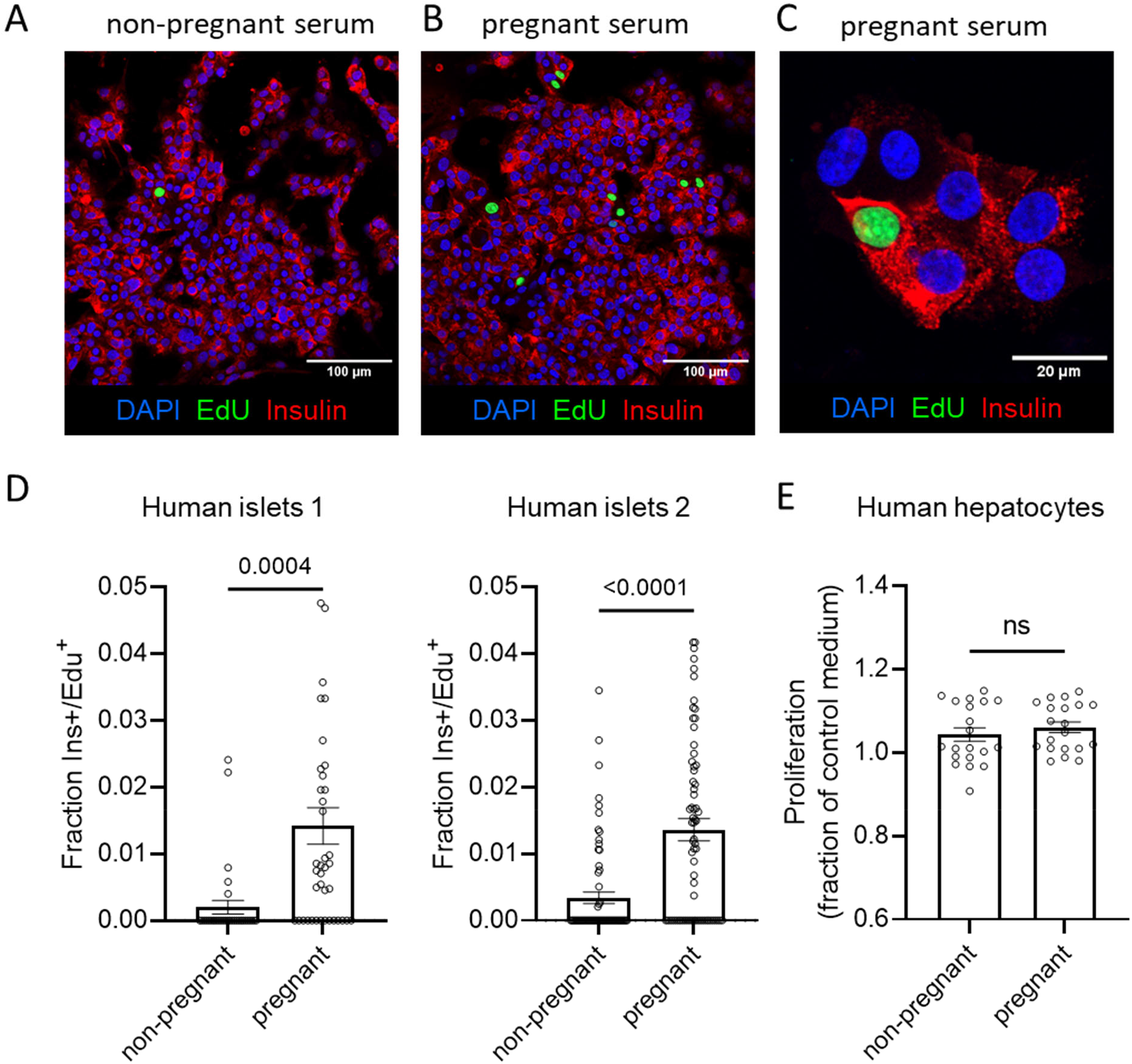
Maternal serum increases proliferation in primary human beta cells but not hepatocytes. (A-C) Monolayer preparations of primary human islets were cultured with 10% pregnant or 10% non-pregnant from pooled donors for two weeks. Proliferation was detected by imaging of EdU incorporation. Right panel shows increased magnification verifying presence in insulin^+^/EdU^+^ beta cells. (D,E) Image quantification of Ins^+^/EdU^+^ cells. Each dot represents EdU positive beta cells for a single image (Islet donor 1: 42 images from 9 wells; Islet donor 2: 70 images from 15 wells). (E) Primary human hepatocytes were cultured with pregnant and non-pregnant human serum. EdU incorporation was measured by EdU microplate assay. Each dot represents separate wells. Groups were assessed for differences by two-tailed t test.

## DISCUSSION

This study is among the first to investigate the effect of human maternal serum on human beta cell proliferation. Factors in the serum of some pregnant donors did enhance human beta cell proliferation and/or insulin secretion *in vitro*, an effect which has also been observed in rat beta cells exposed to maternal human serum [16]. Our study suggests a cell-type specific mechanism(s) at play because primary human hepatocyte proliferation was not affected by serum pregnancy status. Enhanced beta cell function in pregnancy may be a result of increased insulin secretion and not always via beta cell proliferation. This finding is similar to other studies which have found that mouse beta cell mitogens fail to produce comparable human beta cell proliferation but do induce increases in insulin secretion, including growth hormone, prolactin, insulin-like growth factor I, placental lactogen, adiponectin, and serotonin [9, 17-20]. Although much is known about mitogens which enhance proliferation of the mouse islet, loss or blocking of those pathways does not completely block the beta cell response to pregnancy [5, 8, 9].

Therefore, additional signals almost certainly contribute to beta cell proliferation in pregnancy. Also, it is unclear whether mechanisms affecting beta cells in mouse pregnancy are the same in humans.

This study was focused on *in vitro* analyses which does not capture the full complexity of physiological organ systems, especially during pregnancy. Nevertheless, the study focused exclusively on human cells including primary human beta cells and serum from pregnant human donors which provides additional insight into mechanism of human beta cell adaptation to pregnancy. In this study we did not observe a uniform response among donors, suggesting heterogeneity of beta cell mitogenic factors in human pregnancy.

## DATA AND RESOURCE AVAILABILITY

All data points generated or analyzed during this study are included in the published article (and its online supplementary files). Raw data are presented in the supplemental figures. Information on the primary human islets analyzed are available in the IIDP repository, https://iidp.coh.org/ [RRID: SAMN13028024; SAMN32641505]. Pregnant serum donor and human islet donor information is listed in (**Table 1 and Supplementary Tables 1 and 2**). No other applicable resources were generated during the current study.

## Ethical Approvals

This protocol was approved by the Institutional Review Board (IRB) and the University of Florida (UF IRB201902512). Primary human pancreatic islets from deceased nondiabetic donors were obtained from the Integrated Islet Distribution Program (IIDP) at City of Hope. Human pancreas tissues used were approved as non-human by the University of Florida IRB (IRB no. 201701113, 201702860).

## ACKNOWLEDGEMENTS

Human pancreatic islets were provided by the NIDDK-funded Integrated Islet Distribution Program (IIDP) at City of Hope, NIH Grant # 2UC4DK098085 and the JDRF-funded IIDP Islet Award Initiative. Research reported in this publication was supported by the University of Florida Clinical and Translational Science Institute, which is supported in part by the NIH National Center for Advancing Translational Sciences under award number UL1TR001427 and by NIH grant R01DK132387. The content is solely the responsibility of the authors and does not necessarily represent the official views of the National Institutes of Health. We thank the serum donors for agreeing to participate in the study.

## Author Contributions

K.S. and C.R. recruited patients, collected sera, and performed beta cell experiments. A.P. and M.B. performed experiments. D.H. and J.C. setup and cared for beta cell cultures. C.M., C.W., M.A., and R.E. provided critical equipment, reagents, expertise, and support. K.S., M.A., R.E. and E.P. conceived the study. K.S. and E.P. analyzed data and wrote the manuscript. All authors contributed to discussion and reviewed/edited the manuscript.

## Notes

### Competing Interest Statement

The authors have declared no competing interest.

