## Supplementary Information for "Serum from pregnant donors induces human beta cell proliferation and insulin secretion"

**Supplemental Tables and Figures**

| **Donor Number** | **Age** | **Race / Ethnicity** | **BMI (kg/m^2^)** | **GA at delivery^a^** | **Glucose Tolerance** | **Neonatal Sex** | **Birth weight (g)** | **Placental Weight** | **Prolactin (ng/ml)** | **Glucose (mg/dl)** |
| --- | --- | --- | --- | --- | --- | --- | --- | --- | --- | --- |
| 1 | 32 | White | 38 | 39.1 | NGT | Male | 3000 | 602 | 159.85 | 93 |
| 2 | 34 | Hispanic | 53.9 | 37.1 | NGT | Male | 4250 | 900 | 87.86 | 99 |
| 3 | 30 | Black | 35.7 | 37.3 | NGT | Male | 3490 | 660 | 56.06 | 112 |
| 4 | 24 | Black | 45.9 | 39.2 | NGT | Female | 3240 | 670 | 151.35 | 104 |
| 5 | 27 | White | 36.1 | 39.6 | NGT | Female | 3130 | 610 | 399.35 | 94 |
| *6* | 37 | Hispanic | 26.9 | 39 | T2DM | Female | 3360 | 519 | 88.63 | 95 |
| *7* | 31 | White | 36.5 | 38.1 | GDM | Male | 4020 | 500 | 222.9 | 130 |
| *8* | 31 | White | 30.9 | 39 | NGT | Male | 3594 | 499 | 147.04 | 66 |
| *9* | 26 | White | 32.7 | 39.2 | NGT | Male | 3450 | 689 | 245.5 | 52 |
| *10* | 31 | White | 37.6 | 39.3 | GDM | Male | 4220 | 1000 | 177.2 | 109 |
| *11* | 32 | White | 27.4 | 39.2 | NGT | Female | 2920 | 502 | 130.2 | 93 |
| *12* | 29 | White | 29.1 | 39.6 | NGT | Male | 3290 | 470 | 163.53 | 78 |
| *13* | 19 | White | 39.4 | 39.1 | NGT | Male | 1600 | 670 | 107.62 | 86 |
| *14* | 35 | Black | 29.2 | 40 | NGT | Male | 3720 | 640 | 189.78 | 75 |
| *15* | 35 | Other | 32 | 39.2 | NGT | Male | 3950 | 885 | 287.4 | 75 |
| *16* | 22 | White | 32.14 | 39 | NGT | Female | 3110 | 570 | - | - |
| *17* | 32 | White | 52.61 | 39 | NGT | Male | 3550 | 800 | - | - |
| *18* | 37 | White | 36.1 | 39 | NGT | Female | 3310 | 500 | - | - |
| *19* | 31 | White | 34.4 | 39 | NGT | Female | 3780 | 800 | - | - |
| *20* | 43 | Black | 51 | 37.3 | NGT | Male | 3400 | 850 | - | - |

**Supplementary Table 1: Baseline characteristics of all serum donors and their neonates.** Peripheral blood samples from donors 16-20 were hemolyzed and unable to be used for experiments. ^a^GA = Gestational age presented as weeks.days (i.e. 39.6 = 39 weeks and 6 days). NGT = normal glucose tolerant; T1DM = type 1 diabetes mellitus; T2DM = Type 2 diabetes mellitus.

| **RRID** | RRID:SAMN13028024 | RRID:SAMN32641505 |
| --- | --- | --- |
| **age** | 57 | 16 |
| **sex** | female | Male |
| **BMI** | 23.5 | 29.5 |
| **HbA1c** | 4.40% | 5.4% |
| **donor type** | no diabetes | no diabetes |

**Supplementary Table 2: human islet donor characteristics.**

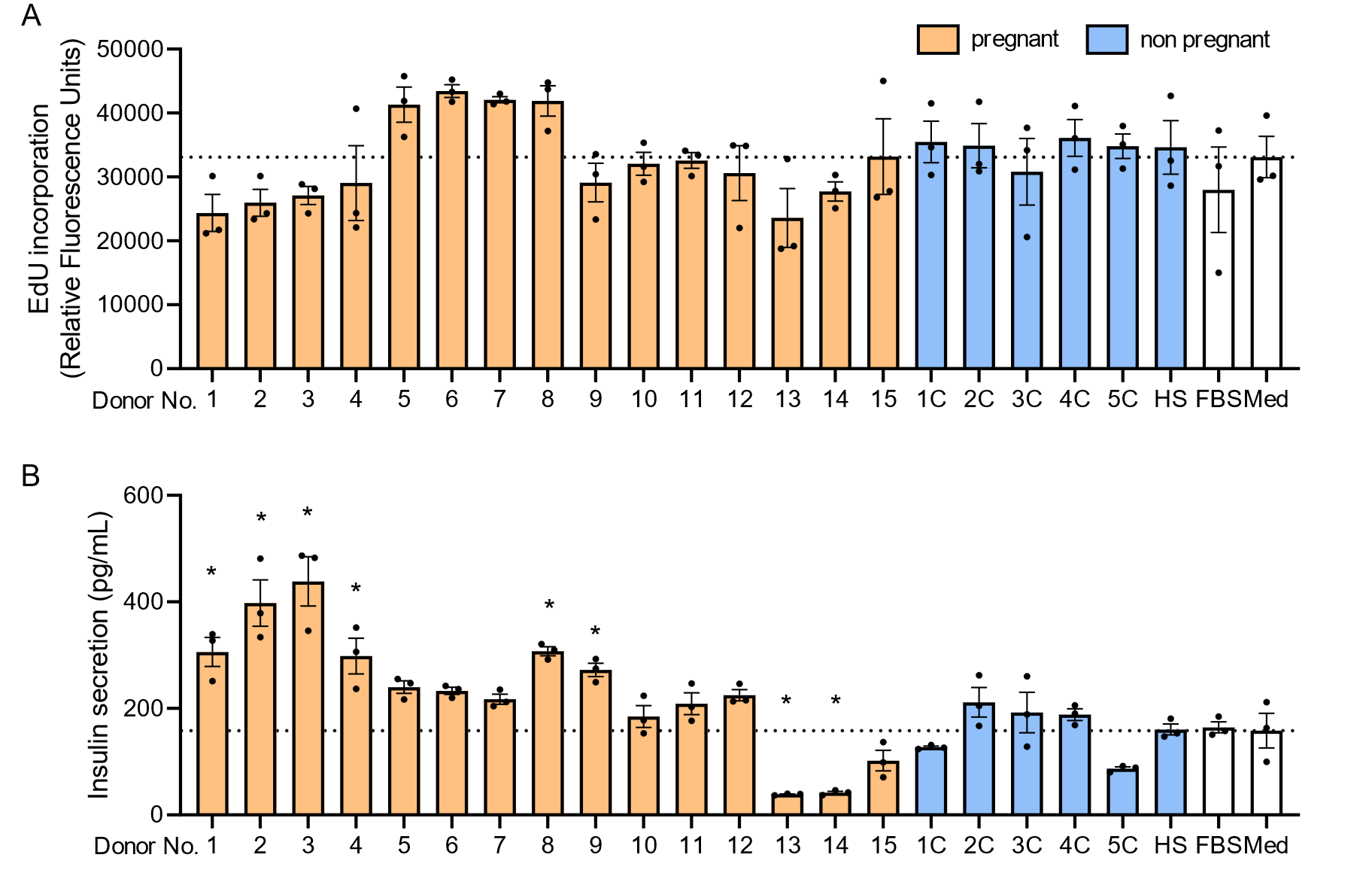

**Supplementary Figure 1. Maternal serum effects on EndoC-βH1 beta cell proliferation and insulin secretion: individual donors’ absolute values.** EndoC-βH1 beta cells were cultured in 96-well plates supplemented with 10% pregnant human serum (Donors 1-15), 10% non-pregnant human serum (Donors 1C-5C), pooled male, non-pregnant human serum (HS), fetal bovine serum (FBS), or control medium (Med) for 7 days. **(A)** Total EndoC-βH1 beta cell proliferation in relative fluorescence units (RFU) measured by EdU incorporation. **(B)** Accumulated EndoC-βH1 beta cell insulin secretion concentration (pg/ml) measures by ELISA kit. Each dot represents experimental replicates (3) for an individual donor. Values are mean ± SEM. *p<0.05 for ANOVA with posthoc pairwise comparisons relative to the mean for control, serum-free media.

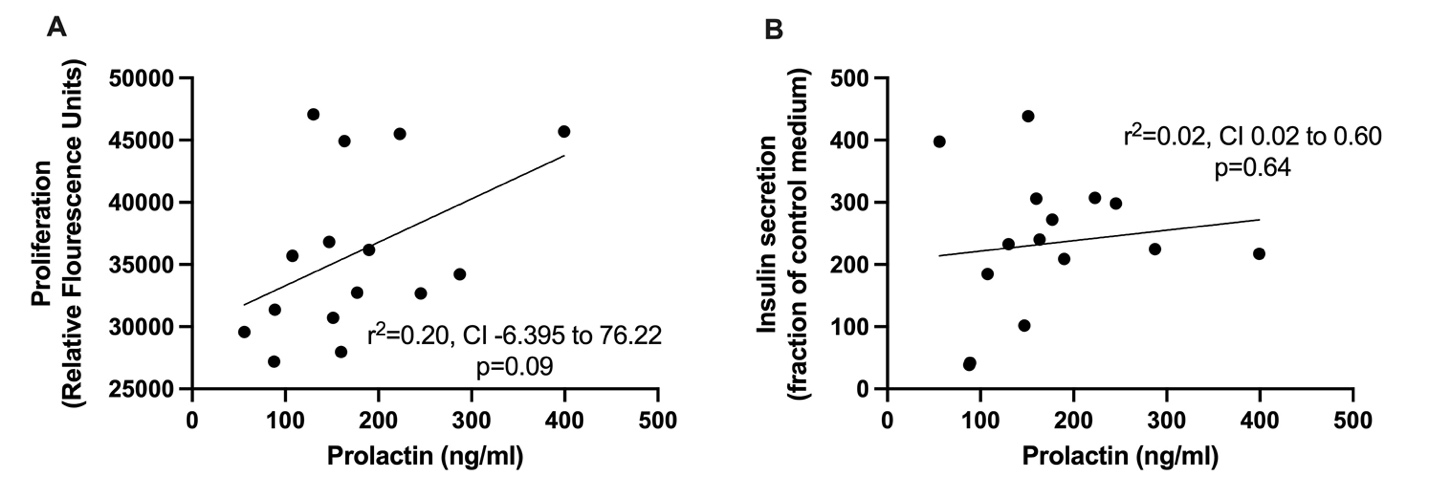

**Supplemental Figure 2. Prolactin concentration compared to EndoC-βH1 beta cell proliferation and insulin secretion.** Prolactin concentration of donors were measured and plotted against proliferation and insulin secretion of EndoC-βH1 beta cells after seven day incubation. **(A)** Relationship between prolactin and EndoC-βH1 beta cell proliferation. **(B)** Relationship between prolactin concentration and EndoC-βH1 beta cell insulin secretion. Spearman’s correlation was used to assess the relationship between prolactin and EndoC-βH1 beta cell proliferation and insulin secretion.
